# FEDERATED MORPHOMETRY FEATURE SELECTION FOR HIPPOCAMPAL MORPHOMETRY ASSOCIATED BETA-AMYLOID AND TAU PATHOLOGY

**DOI:** 10.1101/2021.08.22.457269

**Authors:** Jianfeng Wu, Qunxi Dong, Jie Zhang, Yi Su, Teresa Wu, Richard J. Caselli, Eric M. Reiman, Jieping Ye, Natasha Lepore, Kewei Chen, Paul M. Thompson, Yalin Wang, for the Alzheimer’s Disease Neuroimaging Initiative

## Abstract

Amyloid-β (Aβ) plaques and tau protein tangles in the brain are now widely recognized as the defining hallmarks of Alzheimer’s disease (AD), followed by structural atrophy detectable on brain magnetic resonance imaging (MRI) scans. One of the particular neurodegenerative regions is the hippocampus to which the influence of Aβ/tau on has been one of the research focuses in the AD pathophysiological progress. This work proposes a novel framework, Federated Morphometry Feature Selection (FMFS) model, to examine subtle aspects of hippocampal morphometry that are associated with Aβ/tau burden in the brain, measured using positron emission tomography (PET). FMFS is comprised of hippocampal surface-based feature calculation, patch-based feature selection, federated group LASSO regression, federated screening rule-based stability selection, and region of interest (ROI) identification. FMFS was tested on two ADNI cohorts to understand hippocampal alterations that relate to Aβ/tau depositions. Each cohort included pairs of MRI and PET for AD, mild cognitive impairment (MCI) and cognitively unimpaired (CU) subjects. Experimental results demonstrated that FMFS achieves an 89x speedup compared to other published state-of-the-art methods under five independent hypothetical institutions. In addition, the subiculum and *cornu ammonis* 1 (CA1 subfield) were identified as hippocampal subregions where atrophy is strongly associated with abnormal Aβ/tau. As potential biomarkers for Aβ/tau pathology, the features from the identified ROIs had greater power for predicting cognitive assessment and for survival analysis than five other imaging biomarkers. All the results indicate that FMFS is an efficient and effective tool to reveal associations between Aβ/tau burden and hippocampal morphometry.

## 1. INTRODUCTION

Alzheimer’s disease (AD) is now viewed as a gradual process that begins many years before the onset of detectable clinical symptoms. Measuring brain biomarkers and intervening at preclinical AD stages are believed to improve the probability of therapeutic success (Brookmeyer et al., 2007; Jack et al., 2016; Sperling et al., 2011). Amyloid-β (Aβ) plaques and tau tangles are two specific protein pathological hallmarks of AD and are believed to induce neurodegeneration and structural brain atrophy consequentially observable from volumetric magnetic resonance imaging (MRI) scans (Gordon et al., 2019; Jack et al., 2008; La Joie et al., 2020; Selkoe and Hardy, 2016). Brain Aβ and tau pathology can be measured using positron emission tomography (PET) with amyloid/tau-sensitive radiotracers or by using lumbar puncture to measure these proteins in samples cerebrospinal fluid (CSF). Even so, these invasive and expensive measurements are less attractive to subjects in the preclinical stage, and PET scanning is also not as widely available as MRI.

In the A/T/N system - a recently proposed research framework for understanding the biology of AD - the presence of abnormal levels of Aβ (A in A/T/N) in the brain or cerebrospinal fluid (CSF) is used to define the presence of biological Alzheimer’s disease (Jack et al., 2016). An imbalance between production and clearance of Aβ occurs early in AD and is typically followed by the accumulation of tau (T in A/T/N) protein tangles (another key pathological hallmark of AD) and neurodegeneration (N in A/T/N) detectable on brain MRI scans (Hardy and Selkoe, 2002; Jack et al., 2016; Sperling et al., 2011). Therefore, there has been great interest in developing techniques to associate Aβ and tau deposition with MRI measures (Ansart et al., 2020; Dahl et al., 2021; Ezzati et al., 2020; Petrone et al., 2019; Sun et al., 2020; Ten Kate et al., 2018; Tosun et al., 2021, 2016, 2014, 2013). In the structural MRI, the hippocampus is a primary target region across the spectrum of dementia research from clinically normal to late stages of AD (Cullen et al., 2020; Dong et al., 2019; B. Li et al., 2016; Shi et al., 2011). Cognitively unimpaired individuals with abnormally high Aβ burden have faster progression of hippocampal volume atrophy (Insel et al., 2017; Zhang et al., 2020). Additionally, tau burden in the brain, assessed using PET tracers, also strongly correlates with subsequent hippocampal volume atrophy (La Joie et al., 2020).

However, the influence of Aβ/tau pathology on regional hippocampal atrophy in AD is still not fully understood. A study by (Hanko et al., 2019) examined correlations between 3D hippocampal shape measures and Aβ/tau burden in 42 subjects and reported a significant association between tau burden and atrophy in specific hippocampal subregions (CA1 and the subiculum), but detected no Aβ-associated hippocampal ROIs. Our previous studies observed an association between hippocampal morphometry and Aβ burden on 1,101 subjects (Wu et al., 2021, 2018) and found significant Aβ-associated hippocampal subregions in the CA1 subfield and the subiculum (Wu et al., 2020). Overall, studies of hippocampal ROIs in larger cohorts tend to be more highly powered and reliable.

Integrating data from multi-sites is common practice for large sample sizes and increased statistical power. An important direction of interest in multi-site neuroimaging research is federated learning – which offers an approach to learn from data spread across multiple sites without having to share the raw data directly or to centralize in any one location. In many cases, different institutions may not be readily able to share biomedical research data due to patient privacy concerns, data restrictions based on patient consent or institutional review board (IRB) regulations, and legal complexities; this can present a major obstacle to pooling large scale datasets to discover robust and reproducible signatures of specific brain disorders. To remedy this distributed problem, a large-scale collaborative network, ENIGMA consortium, was built (Thompson et al., 2020). However, most ENIGMA meta-analytic studies currently focus on univariate measures derived from brain MRI, diffusion tensor imaging (DTI), electroencephalogram (EEG), or other data modalities, and relatively few have studied multivariate imaging measures. Federated learning models, such as decentralized independent component analysis (Baker et al., 2015), sparse regression (Plis et al., 2016), and distributed deep learning (Kaissis et al., 2021; Stripelis et al., 2021; Warnat-Herresthal et al., 2021), have made solid progress with leveraging multivariate image features for statistical inferences, allowing iterative computation on remote datasets. Some other recent studies focus on multivariate linear modeling (Silva et al., 2020), federated gradient averaging (Remedios et al., 2020), and unbalanced data for multi-site (Yeganeh et al., 2020). To our knowledge, these methods have not yet been applied to detect multimodal associations in Alzheimer’s disease research, such as finding anatomically abnormal regions on MRI that are associated with Aβ/tau pathology defined using PET.

Here we propose a novel framework, Federated Morphometry Feature Selection (FMFS), to detect the association between hippocampal morphometry markers and Aβ/tau burden. FMFS calculates patch-based surface morphometry features from brain MRI scans of people with AD, mild cognitive impairment (MCI), and cognitively unimpaired (CU) subjects. With our novel federated feature selection method based on group LASSO regression, we apply the proposed framework to assess hippocampal regions of interest (ROIs) associated with Aβ/tau burden (note that by ROIs, we mean subregions and advanced morphometric features on the 3D hippocampal surface, which may have a finer scale than currently defined subregions of the hippocampus).

To test the added value of distributed computing, we also hypothesize that the proposed framework could leverage distributed computational models to improve the statistical power to identify the influence of Aβ/tau pathology on regional hippocampal morphometry. To examine the value of subregional hippocampal features as effective biomarkers of AD progression, we train several regression models with the features from the ROIs to predict the cross-sectional MMSE (mini-mental state exam) score (Folstein et al., 1975) – a very widely used clinical measure of disease severity in AD. In addition, we use a separate dataset to demonstrate our ROIs offer superior performance relative to several other univariate measures in a survival analysis of MCI conversion to AD. Our work generalizes and enriches federated learning research by explicitly selecting (and visualizing) key regional features. By increasing access to information from large-scale imaging datasets and computing efficiency, FMFS may offer an efficient and effective screening tool to reveal the associations between Aβ/tau burden and hippocampal morphology across the dementia spectrum, and the features on ROIs could provide a means for screening individuals prior to more invasive Aβ/tau burden assessments that might determine their eligibility for interventional trials.

## 2. SUBJECTS and METHODS

### 2.1 Subjects

Data used in the preparation of this article were obtained from the Alzheimer’s Disease Neuroimaging Initiative (ADNI) database (adni.loni.usc.edu). The ADNI was launched in 2003 as a public-private partnership led by Principal Investigator Michael W. Weiner, MD. The primary goal of ADNI has been to test whether serial MRI, PET, other biological markers, and clinical and neuropsychological assessments can be combined to measure the progression of MCI and early AD. For up-to-date information, see www.adni-info.org. From the multiple phases of ADNI - ADNI 1, ADNI 2, ADNI GO, and ADNI 3 - we analyzed two sets of scans for the study of Aβ deposition and tau deposition. For the Aβ deposition study, we analyzed a total of 1,127 pairs of images from 1109 subjects (18 of them have two pairs from different visiting dates), including 1,127 T1-weighted MR images and 1,127 florbetapir PET images. Similarly, we obtained 925 pairs from 688 subjects (191 of them have more than one pair from different visiting dates) of MRI scans and AV1451 PET images for the study of tau deposition.

In addition to each brain MRI scan, we also analyze the corresponding Mini-Mental State Exam (MMSE) scores (Folstein et al., 1975). For amyloid PET, we utilize centiloid measures (Navitsky et al., 2018). Operationally, there have been widely accepted efforts to reconcile differences among different amyloid radiotracers using a norming approach called the centiloid scale (Klunk et al., 2015; Rowe et al., 2017). ADNI florbetapir PET data are processed using AVID pipeline (Navitsky et al., 2018), which are converted to the Centiloid scales according to their respective conversion equations (Navitsky et al., 2018; Su et al., 2019). For flortaucipir tau-PET - in a similar fashion to Aβ - tau data are reprocessed using a single pipeline consistent with (Sanchez et al., 2020), so that the standardized uptake value ratio (SUVR) from different ADNI study sites can be analyzed together. In this work, we examine two regional SUVR for tau deposition, corresponding to Braak12, and Braak34 (Baker et al., 2017b, 2017a; Maass et al., 2017; Schöll et al., 2016). **Table 1** shows the demographic information from the two cohorts that we analyzed.

**Table 1.** Demographic information for the participants we studied from the ADNI.

| Cohort | Group | Sex (M/F) | Age | MMSE | Centiloid |  |
| --- | --- | --- | --- | --- | --- | --- |
| Aβ<br>(n=1,127) | AD (n=173) | 98/75 | 75.0±7.8 | 22.7±2.9 | 72.0±40.2 | |
| | MCI (n=516) | 291/225 | 72.6±7.8 | 28.0±1.7 | 42.0±40.7 | |
| | CU (n=438) | 200/238 | 74.5±6.5 | 29.0±1.2 | 24.4±33.3 | |
| Cohort | Group | Sex (M/F) | Age | MMSE | Braak12 | Braak34 |
| tau<br>(n=925) | AD (n=115) | 62/53 | 76.0±8.5 | 22.0±4.5 | 2.39±0.60 | 2.51±0.73 |
| | MCI (n=278) | 158/120 | 74.6±7.9 | 27.9±2.1 | 1.82±0.46 | 1.92±0.46 |
| | CU (n=532) | 210/322 | 73.4±7.1 | 29.1±1.1 | 1.58±0.23 | 1.73±0.21 |

Values are mean ± standard deviation, where applicable.

### 2.2 Proposed pipeline

In this work, we develop a Federated Morphometry Feature Selection (FMFS) model to detect the influence of Aβ and tau deposition on hippocampal shape deformity and to better support the future prediction of AD pathology, as shown in **Figure 1**. In panel (1), each institution first extracts the morphometric features locally. The hippocampal structures are segmented from registered brain MR images, and smoothed hippocampal surfaces are further generated. After the surface parameterization and fluid registration, the hippocampal radial distance (RD) and tensor-based morphometry (TBM) features are calculated at each surface point. Each institution selects patches on each hippocampal surface and reshapes the grouped features (RD or TBM on each patch are one group) of each subject to a vector. Next, in panel (2), taking each Aβ/tau measurement as the dependent variable, the institutions perform the federated feature selection model on these patches of features to generate local hippocampal regions of interest (ROIs) for each Aβ/tau measurement.

**Figure 1.**
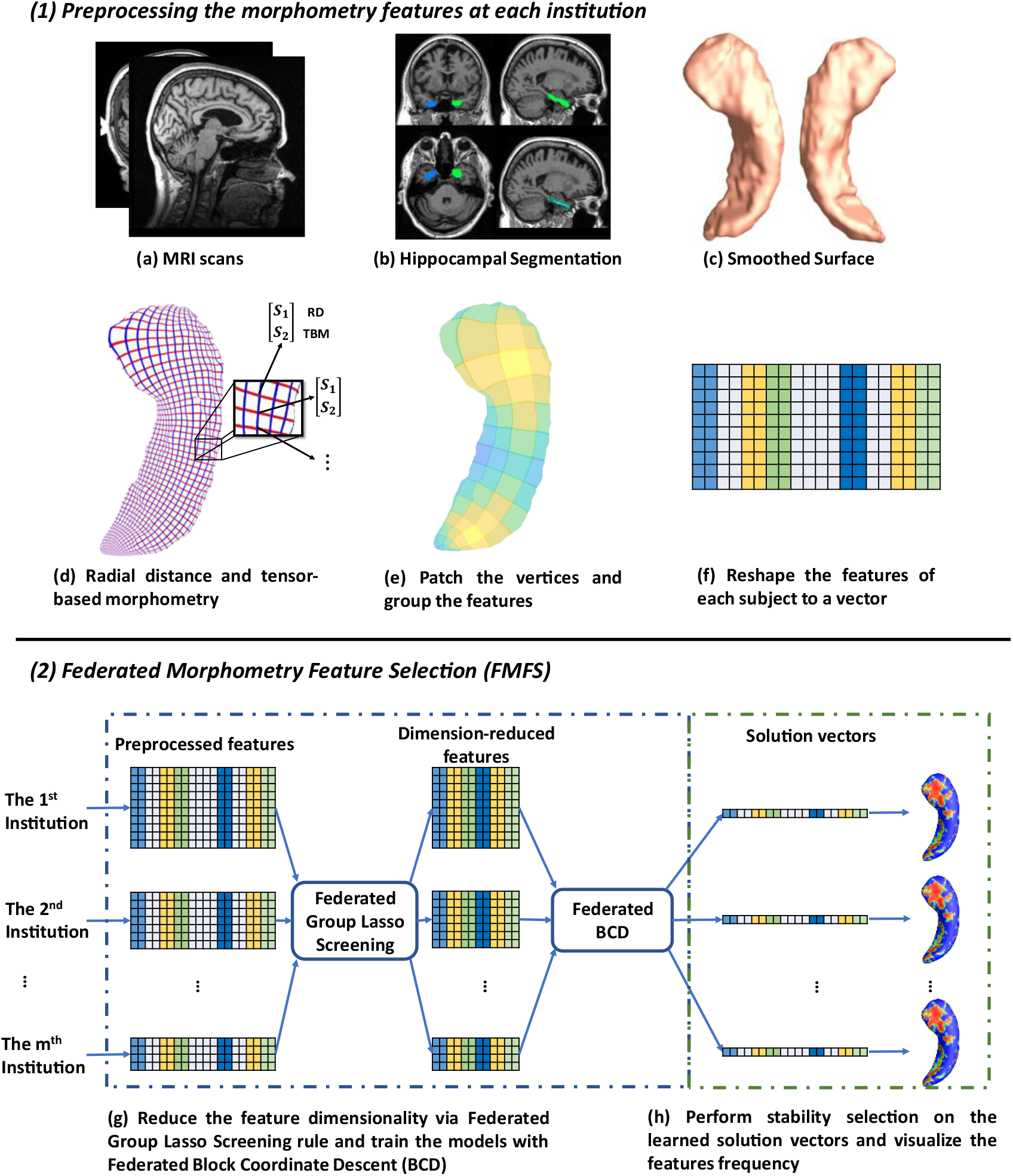
System pipeline. Panel (1) shows the steps for each institution to extract morphometric features locally. The hippocampal structures are segmented from registered brain MR images and smoothed hippocampal surfaces are then generated. After the surface parameterization and fluid registration, the hippocampal radial distance (RD) and tensor-based morphometry (TBM) features are calculated at each surface point. Each institution selects patches on each hippocampal surface and reshapes the grouped features of each subject into a vector. Next, in panel (2), taking Aβ/tau measurements as dependent variables, the institutions perform the federated feature selection model on these patches of features to generate hippocampal local regions of interest (ROIs) for each Aβ/tau measurement.

#### 2.2.1 Image processing

Using the FIRST algorithm from the FMRIB Software Library (FSL), hippocampal structures are segmented in the MNI152 standard space (Paquette et al., 2017; Patenaude et al., 2011) (**Figure 1** (b)). Surface meshes are constructed based on the hippocampal segmentations with the marching cubes algorithm (Lorensen and Cline, 1987) and a topology-preserving level set method (Han et al., 2003). Then, to reduce the noise from MR image scanning and to overcome partial volume effects, surface smoothing is applied consistently to all surfaces. Our surface smoothing process consists of mesh simplification using progressive meshes (Hoppe, 1996) and mesh refinement by the Loop subdivision surface method (Loop, 1987) (**Figure 1** (c)). Similar procedures adopted in a number of our prior studies (Colom et al., 2013; Luders et al., 2013; Monje et al., 2013; Shi et al., 2015, 2013b, 2013a; Wang et al., 2012, 2010) show that the smoothed meshes are accurate approximations to the original surfaces, with a higher signal-to-noise ratio (SNR).

Using the holomorphic flow segmentation method (Wang et al., 2007), each hippocampal surface is parameterized with refined triangular meshes, and the parameterized surfaces are then registered to a standard rectangular grid template using a surface fluid registration algorithm (Shi et al., 2013a). After parameterization and registration, we establish a one-to-one correspondence map between hippocampal surfaces. Each surface has the same number of vertices (150 × 100). As illustrated in **Figure 1** (d), the intersection of the red curve and the blue curve is a surface vertex, and at each vertex, we adopt two kinds of morphometry features, the radial distance (RD) (Pizer et al., 1999; Thompson et al., 2004) and measures derived from surface tensor-based morphometry (TBM) (Chung et al., 2008; Davatzikos, 1996; Thompson et al., 2000; Woods, 2003). The RD (a scalar at each vertex) represents the thickness of the shape at each vertex relative to the medial axis; this primarily reflects surface differences along the surface normal directions. The medial axis is determined by the geometric center of the isoparametric curve on the computed conformal grid (Wang et al., 2011). The axis is perpendicular to the isoparametric curve, so the thickness can be easily calculated as the Euclidean distance between the core and the vertex on the curve. TBM examines the Jacobian matrix *J* of the deformation map that registers the surface to a template surface (Shi et al., 2013a). For TBM, 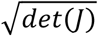 was computed at each vertex, and this value reflects how the surface area changed around the vertex (expansion or atrophy). Additionally, we used the heat kernel smoothing algorithm (Chung et al., 2005; Shi et al., 2015) to refine the surface features. Since the surface of the hippocampi in each brain hemisphere has 15,000 vertices and each vertex has one RD and one TBM, the final feature dimensionality of both hippocampi combined, for each subject, is 60,000 ((15000 + 15000) × 2).

Finally, on each hippocampal surface (100 × 150 vertices), we uniformly selected 2,500 patches of size 2 × 3, and RD and TBM in one patch were considered as a group of features, respectively (**Figure 1** (e)). We selected this patch size of 2 × 3 to increase the robustness of the feature selection model, but also because it does not have an adverse impact on the feature visualization. The grouped features for each subject are reshaped to a vector (**Figure 1** (f)) and will be further processed with our Federated Morphometry Feature Selection (FMFS) model.

#### 2.2.2 Federated Group Lasso Regression

Group LASSO (Yuan and Lin, 2006) is a widely-used technique for group-wise feature selection in high dimensional data. A group-LASSO linear regression has the following optimizing problem:

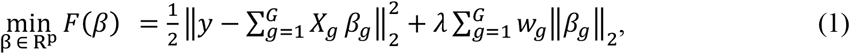

where *X_g_* ∈ R^*N*×*p_g_*^ is the feature matrix, and *y* denotes the *N* dimensional response vector. Group LASSO divides the original feature matrix *X* ∈ R^*N*×*p*^ into *G* different groups, where *X_g_* represents the features in *g*th group and *w_g_* is the weight for this group. After solving the group LASSO problem, we get the *G* solution vectors, *β*_1_,*β*_2_,…,*β_G_*. The dimensionality of each group, *p_g_*, can be arbitrary and the whole solution vector *β* is [*β*_1_,*β*_2_,…,*β_G_*] ∈ R^*p*^. Additionally, *λ* is a positive regularization parameter to control the sparsity of the solution vector, and *w_g_* is the weight for *g*th group of features.

There are many optimization methods to solve the group LASSO problem; block coordinate descent (BCD) (Qin et al., 2013) is one of the most efficient. Instead of updating all the variables at the same time, BCD only updates one or several blocks of variables at each epoch. Therefore, for the group LASSO problem, it can optimize one group of variables while keeping the other ones fixed. Based on this idea, we proposed a federated block coordinate descent (FBCD) to solve our problem.

Li et al., (2016) proposed an optimization model, the local query model (LQM), which preserves the data privacy at each institution. We assume that there are *I* institutions, and each of them owns a private data set (*X^i^*, *y^i^*). We can reformulate the problem (1) as

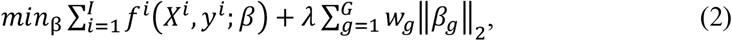

where 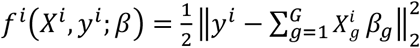 is the least square loss of the *i*th institution. We then have the global gradient,

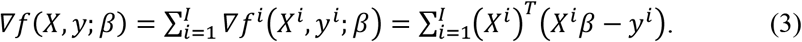

Each of the local institutions calculates its own gradient locally and uploads it to the master server. The latter is used only to determine the gradient with respect to *X*, by adding all ∇*f^i^*(*X^i^, y^i^; β*). It then assigns the global update gradient ∇*f*(*X, y; β*) back to all the local institutions to compute β. Our proposed Federated Block Coordinate Descent (FBCD) method is outlined in **Alg. 1**.

##### Algorithm 1. Federated Block Coordinate Descent (FBCD)

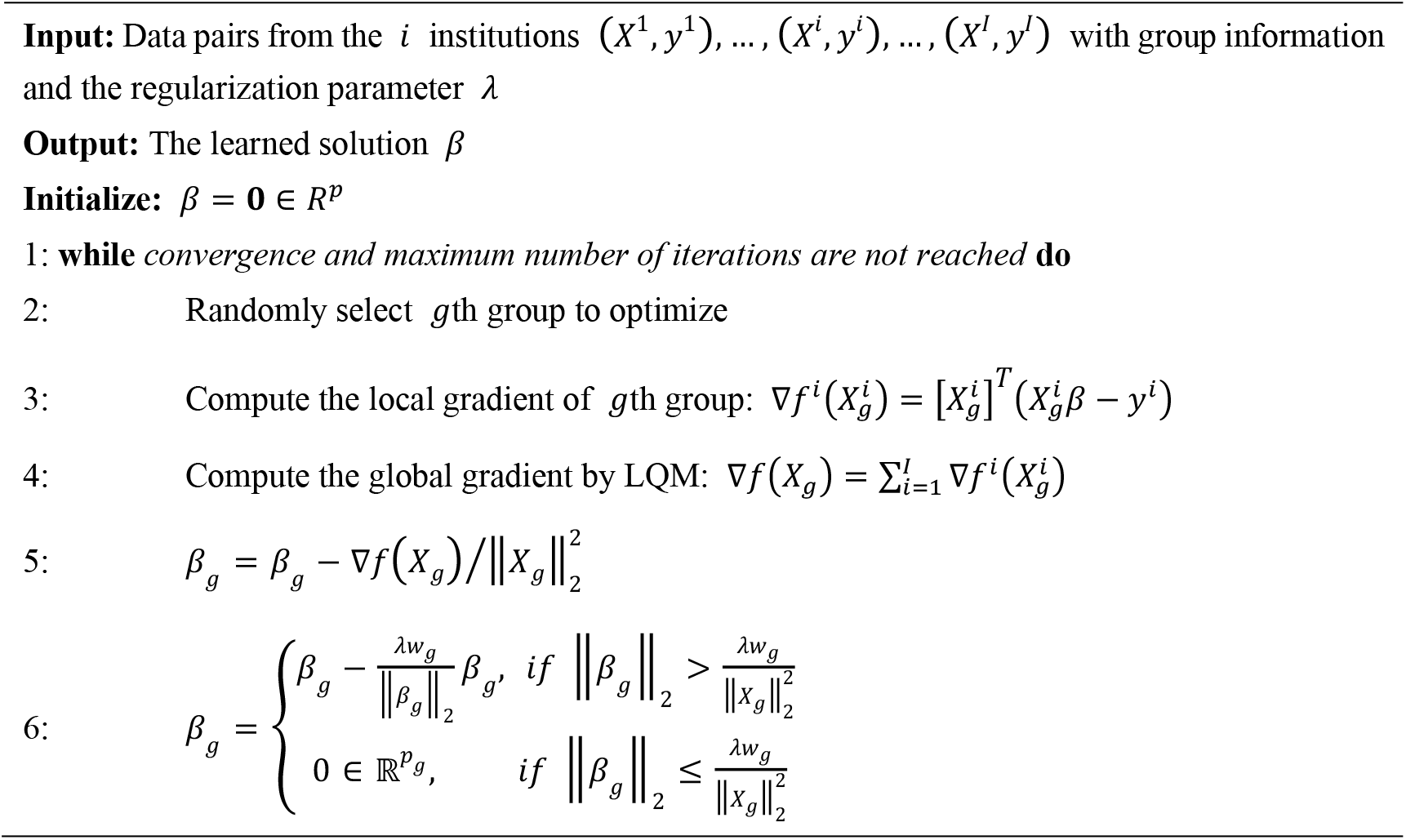

#### 2.2.3 Federated Screening for Group Lasso

Finding the optimal value for the regularization parameter *λ* is a common problem in LASSO techniques. The most frequently used methods, such as cross-validation and stability selection, solve it by trying a sequence of regularization parameters, *λ*_1_ >… > *λ_κ_*; this can be very time-consuming. Instead, the enhanced dual polytope projection rule (EDPP) (Wang et al., 2015) achieved a 200x speedup on the cross-validation in real-world applications, by using information derived from the solution of the previously tried regularization parameter. For the group LASSO problem, the *g*th group of features *X_g_* can be discarded if it satisfies the screening rule, ∥*J_g_*∥ ≤ *w_g_*(2*λ* - *λ_max_*) where 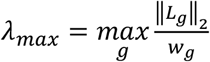 and *J_g_* and *L_g_* are the elements of *J* and *L* defined in **Alg. 2**. The screening rule is based on the uniqueness and non-expansiveness of the optimal dual solution, because the feasible set in the dual space is a convex and closed polytope. More information on EDPP may be found at the following GitHub: http://dpc-screening.github.io/glasso.html

Following the screening rule, we further propose a federated screening rule for group LASSO, named federated dual polytope projection for group LASSO (FDPP-GL), to rapidly locate the inactive features in a distributed manner while preserving data privacy at each institution (**Figure 1** (g)). We summarize the method in **Alg. 2**. In the algorithm, we estimate the maximum regularization parameter, *λ_max_*. The input sequence of parameters, *λ*_1_,*λ*_2_,…,*λ_κ_*, should be no greater than *λ_max_*. Based on the solutions with the sequence of regularization parameters, we can then perform stability selection (Meinshausen and Bühlmann, 2010) to select significant features that are most related to the corresponding *y* (**Figure 1** (h)).

##### Algorithm 2. Federated Dual Polytope Projection for Group Lasso (FDPP-GL)

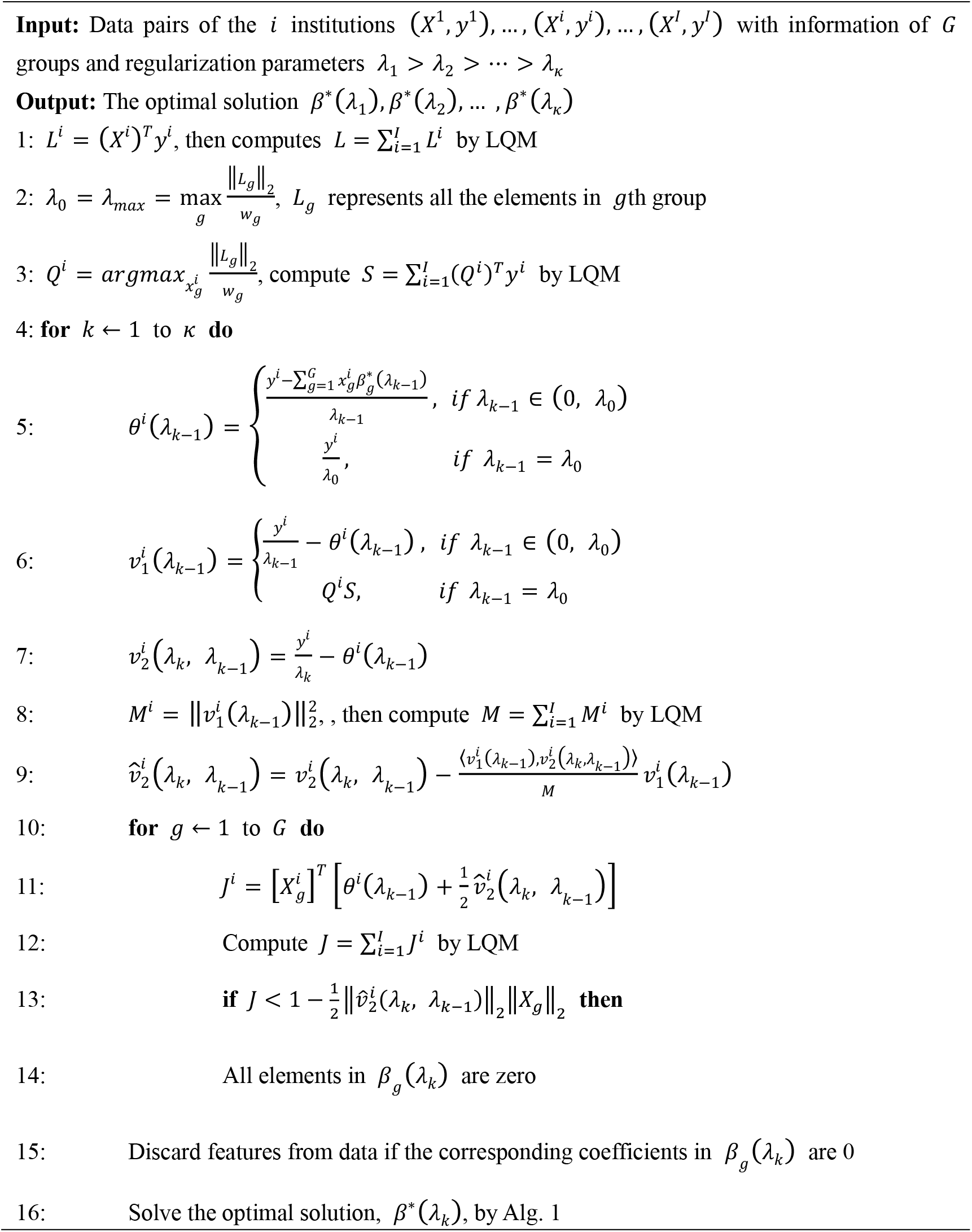

#### 2.2.4 Morphometry Feature Selection and Visualization

We carry out the proposed federated group LASSO method to measure how significantly the patches of features are related to the response *y*. Given a decreasing sequence of regularization parameters, *λ*_1_,…,*λ_κ_*, we learn a set of corresponding models, *β*(*λ*_1_),…,*β*(*λ_κ_*). We perform stability selection by counting the frequency of nonzero entries in the learned models and visualize the frequency on the surface. The counted frequency on each vertex is normalized to 1 to 100 and then mapped to a color bar. For better visualization, we smooth the values on each surface with a 2 × 3 averaging filter. The warmer colored areas have a higher frequency value. In other words, these areas have more significant associations with *y*.

### 2.3 Performance Evaluation Protocol

To further validate whether these identified hippocampal ROIs are related to Aβ/tau deposition, we used RD and TBM features of these ROIs to predict MMSE scores based on random forest, multilayer perceptron (MLP), and LASSO regression models. 10-fold cross-validation was adopted to evaluate the performance of the models, and root mean squared error (RMSE) was used for measuring the prediction accuracy. Meanwhile, we also compared the prediction results of ROI-related features with the results of the whole hippocampal features and Aβ/tau measures.

We also tested the computing efficiency with the 1,127 subjects for the study of Aβ. Firstly, we randomly assign the 1,127 subjects to five institutions, of which each has almost the same number of subjects and one computation node. After uniformly selecting 100 regularization parameters from 1.0 to 0.1, we performed stability selection with our proposed framework, FMFS, FBCD (FMFS without the screening rule), as well as the state-of-the-art distributed alternating direction method of multipliers (DADMM) (Boyd et al., 2011). Besides this, we also repeated the same experiments with different dimensionality of features by randomly down-sampling and up-sampling the original features.

## 3. RESULTS

### 3.1 Efficiency Evaluation

A significant innovation of FMFS is that we introduce a screening rule during the group LASSO feature selection stage, which highly improves the computation speed compared to the distributed alternating direction method of multipliers algorithm (DADMM) (Boyd et al., 2011). Besides, we also tested the running time of FBCD in our federated framework without the screening rule.

We simulated the distributed condition on a cluster with several conventional x86 nodes, of which each contains two Intel Xeon E5-2680 v4 CPUs running at 2.40 GHz. Each parallel computing node has a full-speed Omni-Path connection to every other node in its partition. 1,127 subjects for the Aβ deposition study were randomly assigned to five simulated institutions, each of which has almost the same number of subjects and one computation node. We uniformly selected 100 regularization parameters from 1.0 to 0.1 and ran all three methods with the same experimental set-up. Under different morphometry feature resolutions (where we randomly down-sampled or up-sampled the dimension of the features), our FMFS method achieved a speedup of 62-fold, 80-fold, 86-fold, and 89-fold, compared to DADMM as shown in **Figure 2**. For FBCD, our FMFS has a speedup of 54-fold, 72-fold, 80-fold, and 86-fold.

**Figure 2.**
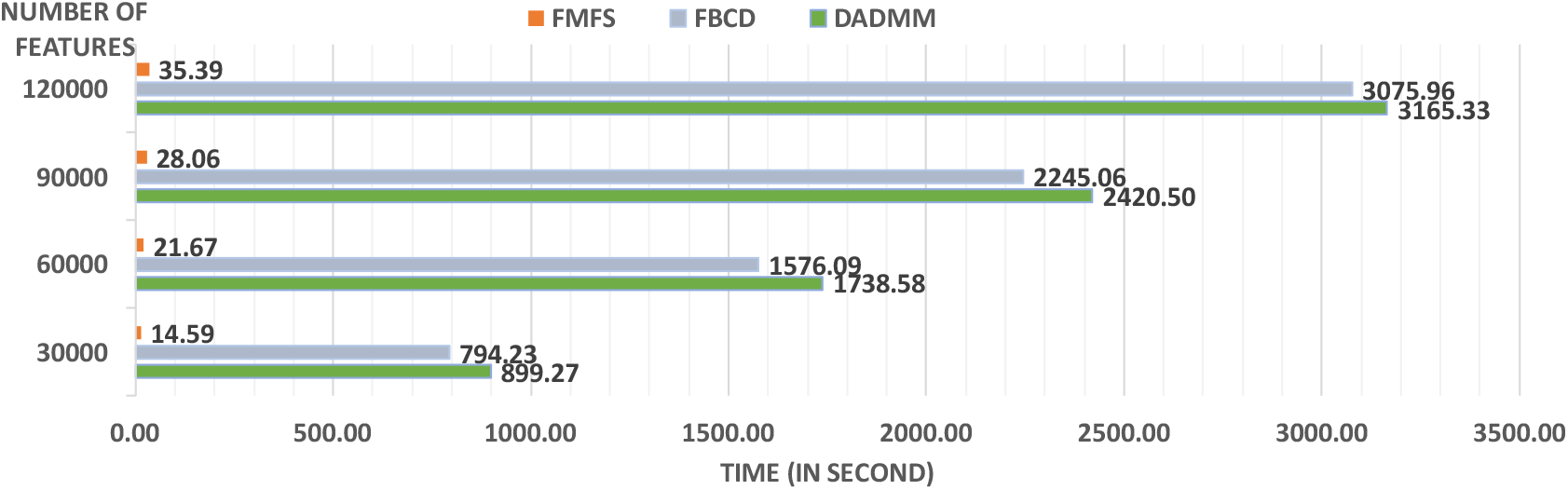
Comparison analysis of computation efficiency. For the morphometry features with different resolutions, our framework achieves a speedup of 62-fold, 80-fold, 86-fold, and 89-fold, compared to DADMM. For FBCD, our FMFS has a speedup of 54-fold, 72-fold, 80-fold, and 86-fold.

### 3.2 Aβ and Tau Associated Hippocampal Morphometry

We employed stability selection with our FMFS model to select the ROIs (subregions of the hippocampal surfaces) related to Aβ and tau. We respectively standardize the two types of input features, RD and TBM, for each subject, using Z-scores, and adopt the centiloid value as the measure of Aβ burden and Braak12 and Braak34 measures for tau deposition. Since the regularization parameters can control the sparsity of the solution vector and further influence the area of the ROIs, we uniformly generated 100 regularization parameters between 0.01 to 0.001, which may lead to a reasonable size for the selected ROIs. After training the model, we got 100 solution vectors, of which the dimensionality is 60,000, since each of the left and right hippocampal surfaces contains 15,000 vertices, and each vertex has two features. Then, we performed stability selection by counting the nonzero entries for RD and TBM on the same vertex. The counted frequency on each vertex was normalized to 0 to 100 and then mapped to a color bar, as shown in **Figure 3-5**. For better visualization, we smoothed the values on each surface by a 2 × 3 averaging filter. The warmer color areas have a higher frequency value and have stronger associations with the responses, i.e., brain global Aβ and tau burden.

**Figure 3.**
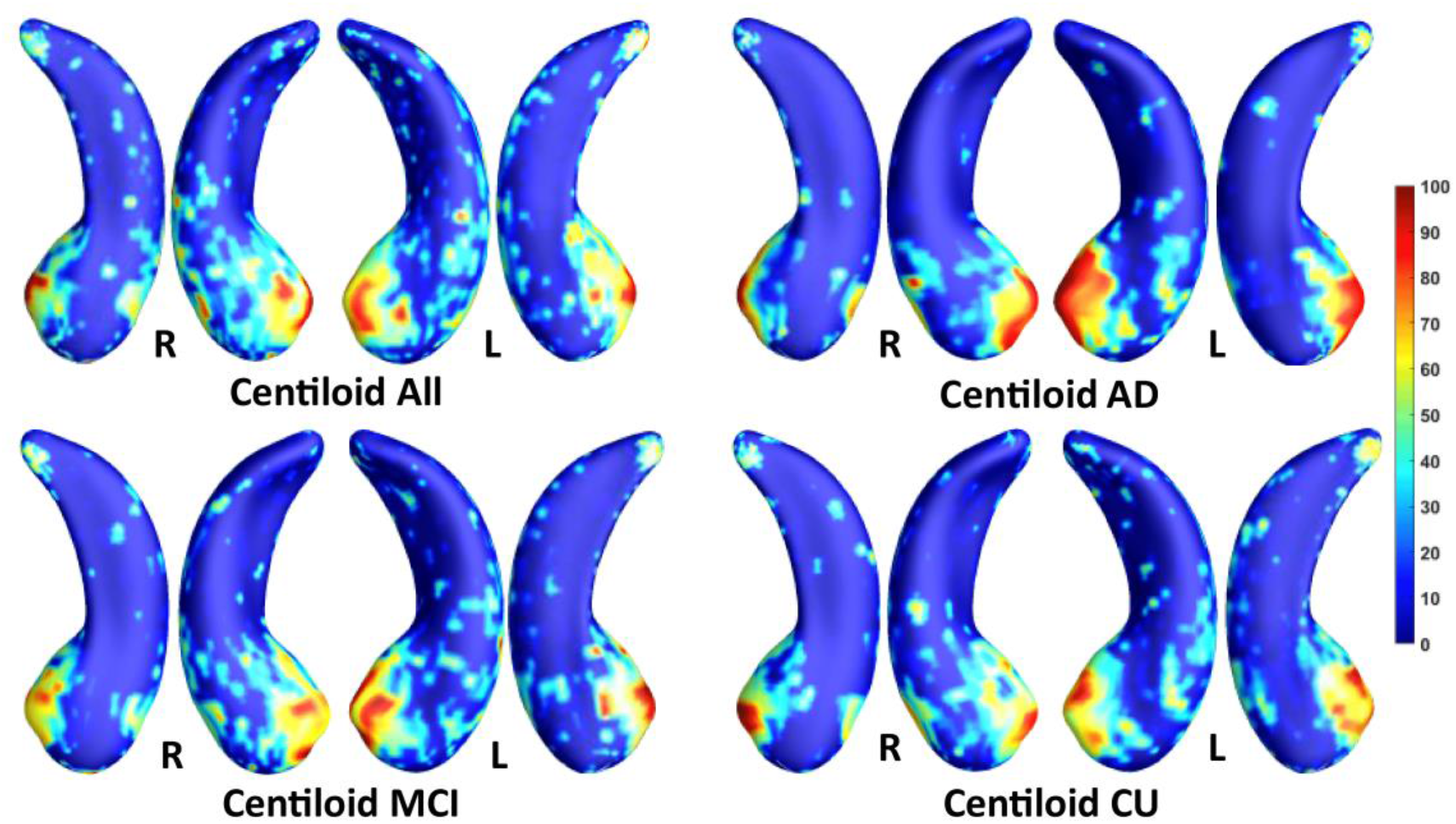
Visualization of ROIs associated with centiloid (Aβ measurement). The top left figure shows the results for all subjects. The top right is for AD patients. The bottom two figures are for participants with MCI and for the CU group.

**Figure 4.**
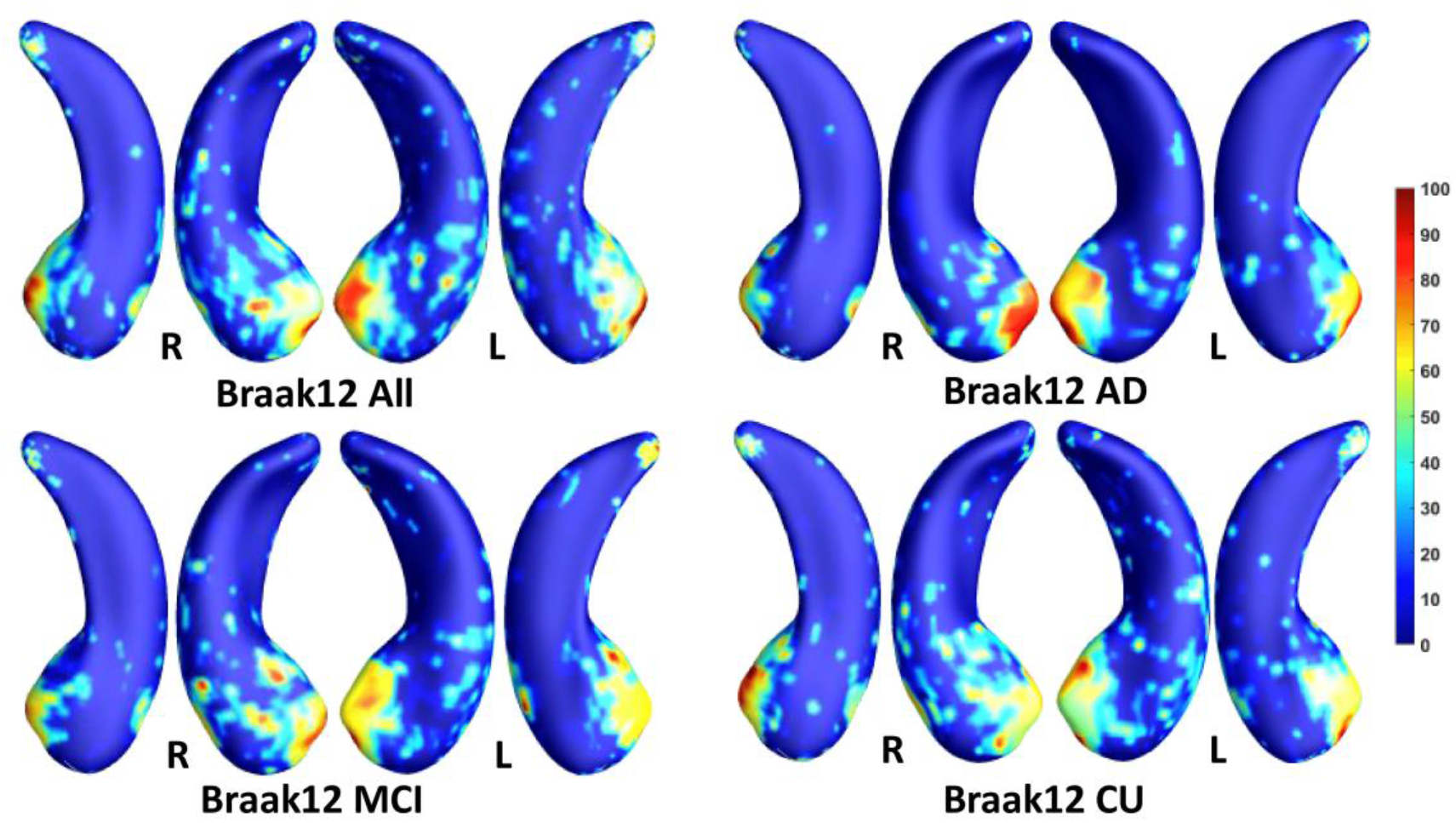
Visualization of ROIs associated with Braak12 (tau measurement). The top left figure shows the results for all subjects. The top right is for AD patients. The bottom two figures are for participants with MCI and for the CU group.

**Figure 5.**
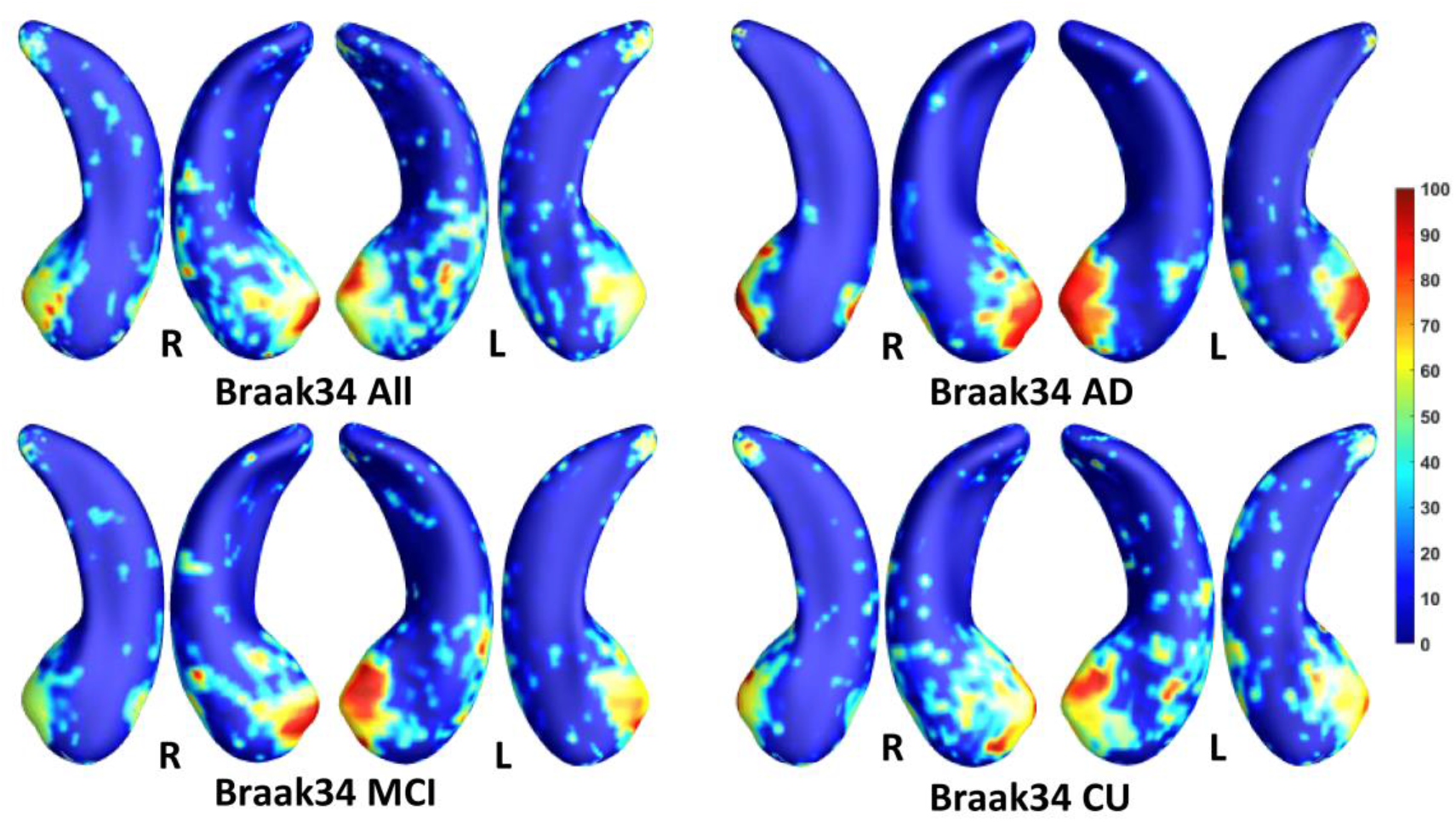
Visualization of ROIs associated with Braak34 (tau measurement). The top left figure shows the results for all subjects. The top right is for AD patients. The bottom two figures are for participants with MCI and for the CU group.

In this experiment, we first ran the proposed model on the cohorts for Aβ and tau deposition. As illustrated in the top left picture of **Figure 3-5**, the morphometric abnormalities mainly happen in specific hippocampal subregions, namely the subiculum and CA1. Additionally, we separately studied the ROIs for groups of CU, MCI, and AD subjects. As shown in the rest of the three panels in **Figure 3-5**, the morphometric associations are strongest in the subiculum and CA1 at the early stages; but with the progression of AD, the distortions are more focal in subiculum. Specifically, the results for CU subjects are shown in the top right panel of each figure, where the warmer colored regions are widespread in both the subiculum and CA1 areas. However, in the results for the AD group, the warmer colored regions mainly focus on the area of the subiculum.

### 3.3 Predicting MMSE Scores based on Hippocampal ROIs

In the model of Jack et al. (2016), an abnormal level of Aβ and tau deposition tends to occur earlier than abnormal cognitive decline can be detected. In this experiment, we further validated the ROIs selected by our proposed model in terms of their prediction accuracy for MMSE score in cohorts where Aβ and tau deposition were measured separately. After performing stability selection, we were able to rank the vertices related to each measurement of Aβ/tau deposition. We selected the 3,000 features from the 1,500 top-ranked vertices for each subject (1,500 RD and 1,500 TBM). Then, we used these features to predict the MMSE score as described in **Section 2.3**. For a fair comparison, we also selected 3,000 features from 1,500 randomly selected vertices for each subject and used them as features representing differences across the entire hippocampus. Moreover, we also leveraged the measurements for Aβ or tau deposition to predict MMSE. In **Table 2**, the top four rows indicate the results for Aβ deposition, and the rest of the rows are for different measurements of tau deposition. These results demonstrate that the features in the ROIs selected by our model can always have a stronger predictive power and predict the MMSE score better than the measurements of Aβ and tau deposition. Our work validated the AD progression model and may provide unique insights for an accurate estimation of clinical disease burden.

**Table 2.** RMSEs for predicting MMSE based on various kinds of biomarkers and models.

| Aβ associated | Random forest | MLP | LASSO |
| --- | --- | --- | --- |
| Hippocampal ROIs | **2.58** | **3.00** | **2.59** |
| Whole Hippocampal | 2.79 | 3.96 | 2.79 |
| Centiloid | 3.15 | 4.01 | 2.85 |
| Braak12 associated | Random forest | MLP | LASSO |
| Hippocampal ROIs | **2.61** | **3.20** | **2.90** |
| Whole Hippocampal | 3.11 | 4.24 | 3.00 |
| Braak12 | 3.03 | 4.98 | 3.03 |
| Braak34 associated | Random forest | MLP | LASSO |
| Hippocampal ROIs | **2.62** | **3.26** | **2.86** |
| Whole Hippocampal | 3.09 | 4.16 | 3.02 |
| Braak34 | 2.81 | 4.94 | 3.02 |

### 3.4 Predicting Clinical Decline in Participants with MCI

In this experiment, we evaluated the performance of our features on the ROI in survival analysis by using 118 MCI participants’ data from a separate dataset (Wang et al., 2021) from ADNI (**Table 3**), including 63 MCI converters, who converted to probable AD in the next six years, and 55 MCI non-converters. Similar to **Section 3.3**, we also chose 1,500 RD and 1500 TBM from the four ROIs (Aβ, Braak12, Braak34) and 3000 features from 1500 random-selected vertices on the whole hippocampal surface to predict the conversion rates from MCI to AD, respectively. For comparison, we also performed the same experiment with the surface area and volume of the hippocampus. The hippocampal volume and surface area were calculated with the smoothed hippocampal structures after linearly registered to the MNI imaging space (Patenaude et al., 2011; Shi et al., 2013a), and the sum of the bilateral hippocampal volume and the sum of the bilateral hippocampal surface area for each subject were used for this experiment.

**Table 3.** Demographic information for participants with MCI.

| Group | Sex (M/F) | Age | MMSE |
| --- | --- | --- | --- |
| MCI converter (n=63) | 42/21 | 75.2±7.0 | 26.7±1.7 |
| MCI non-converter (n=55) | 38/17 | 74.7±7.8 | 27.7±1.4 |

Values are mean ± standard deviation, where applicable.

To fit the univariate Cox model, we converted the features on ROIs to a single value for each subject. First, as the features on the ROIs should have stronger predictive power, we used the frequency on each vertex as a weight to multiply the RD and TBM on the vertex. And then, we respectively summed up the weighted RD and weighted TBM on the ROIs for each subject. The value for RD and the value for TBM were further reduced to a scalar with principal components analysis (PCA). PCA is an unsupervised model to reduce the dimensionality of the data while minimizing information loss. It creates new uncorrelated features which maximize the variance successively. For the randomly selected features on the whole hippocampal surface, the RD and TBM were directly summed up without multiplying the frequency and reduced to a single value with PCA.

Then, the optimal cutoffs for these measurements were determined with the maximum sensitivity and specificity for distinguishing MCI converters and non-converters based on Receiver Operating Characteristic (ROC) analysis (Robin et al., 2011). The ROC curves are illustrated in **Figure 6**, and the AUC, 95% confidence interval (CI) of AUC, and the optimal cutoffs are shown in **Table 4**.

**Figure 6.**
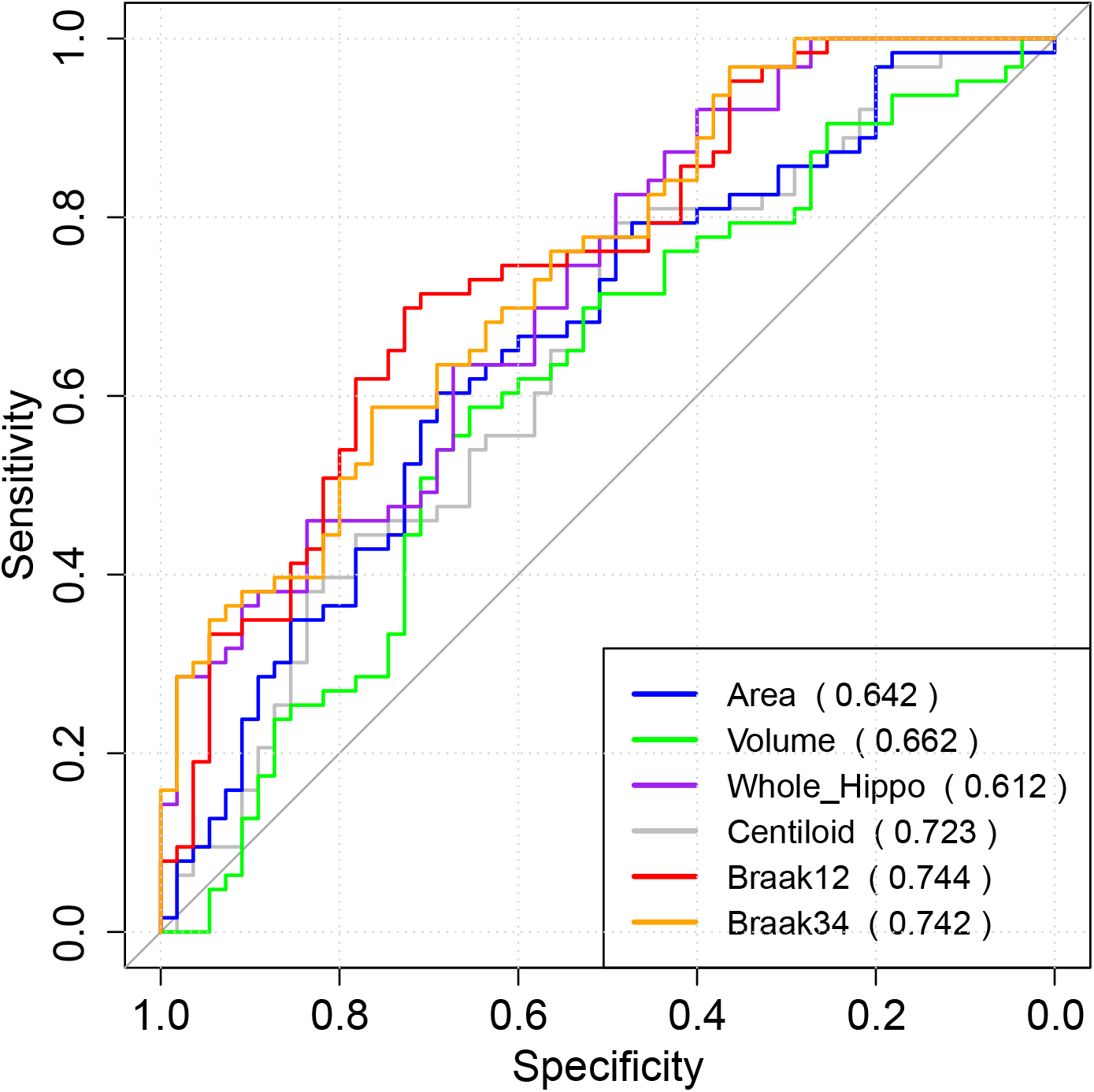
The ROC analysis results for hippocampal surface area, volume, the whole hippocampal feature, and the features on ROIs associated with Aβ, Braak12, and Braak34. The AUC for each measurement is shown in parentheses.

**Figure 7.**
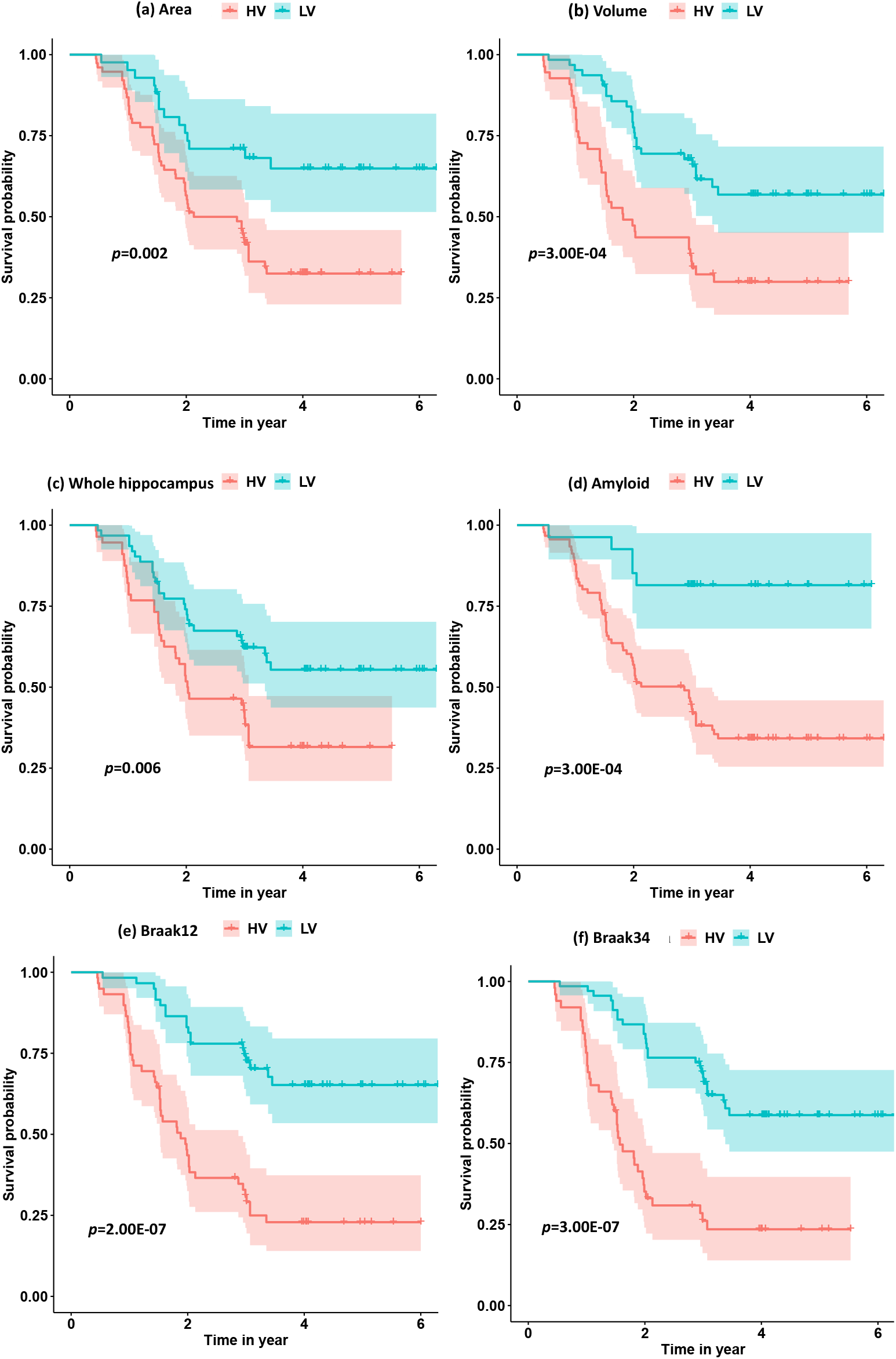
The survival probability analysis for progression to AD in MCI patients based on hippocampal surface area, volume, the whole hippocampal features, and the features on ROIs related to Aβ, Braak12 and Braak34. The *p*-values are from the log-rank test.

**Table 4.** AUC for ROC analysis, optimal cutoffs, and estimated hazards ratios (HRs) for conversion to AD in MCI patients with high-value and low-value biomarkers based on a univariate Cox model.

| Measurements | AUC (95% CI) | Cutoff | $\beta$ | HR (95% CI) | *p*-value |
| --- | --- | --- | --- | --- | --- |
| Area | 0.64 (0.54, 0.74) | 8037.8 | 0.40 | 2.5 (2.3, 3.2) | 0.001 |
| Volume | 0.66 (0.56, 0.76) | 7814.9 | 0.41 | 2.5 (2.2, 3.1) | 4.00E-04 |
| Whole_hippo | 0.61 (0.51, 0.72) | 1.7 | 0.50 | 2.0 (1.7, 2.8) | 0.007 |
| Centiloid | 0.72 (0.63, 0.81) | 13.3 | 0.21 | 4.7 (4.6, 5.2) | 4.00E-05 |
| Braak12 | 0.74 (0.66, 0.83) | -3.0 | 0.26 | 3.8 (3.6, 4.3) | 4.00E-07 |
| Braak34 | 0.74 (0.65, 0.83) | -7.6 | 0.29 | 3.5 (3.3, 4.0) | 1.00E-06 |

With the optimal cutoffs, we could divide the whole cohort into two groups with different measurements. For example, the subjects with hippocampal volume higher than 7814.9 mm^3^ were assigned to a high-value (HV) group, and the rest were into a low-value (LV) group. As expected, AD may decrease the hippocampal volume as well as the other measurements. Next, we fitted a Cox proportional hazard model (Moore, 2008) with the six measurements separately, and the regression beta coefficients (β), the hazard ratios (HRs), and statistical significance (*p*-values) are shown in **Table 4**.

Moreover, we calculated the survival probabilities for conversion to AD in the HV group and the LV group by fitting Kaplan-Meier curves. The survival probabilities of the subjects based on hippocampal surface area, volume, the whole hippocampal features, and the features on ROIs related to Aβ, Braak12, and Braak34 are shown in **Figure 8**. Each color represents the survival curve and 95% CI of one group. Here a log-rank test was used to compare the survival group differences based on a *χ*^2^ test, and the *p*-values are illustrated in each plot. A result with a *p*-value < 0.05 indicates that the two groups are significantly different in terms of survival time. The features from our selected ROIs tended to always yield stronger significant results than the hippocampal surface area, volume, and the whole hippocampal features.

## 4. DISCUSSION

This work proposes a novel framework, FMFS, to efficiently detect Aβ/tau associated hippocampal morphometry markers at different clinically defined stages of AD. The first contribution of this work is that our proposed FMFS model shows excellent computational efficiency compared to similar federated learning models, with a speedup of up to 89-fold. Our work may help accelerate large-scale neuroimaging computations over various disparate, remote data sources without requiring the transfer of any individual data to a centralized location. The second contribution is that the FMFS is an effective tool to select and visualize the brain imaging feature data. In our previous studies (Stonnington et al., 2021; Zhang et al., 2021a, 2021b), the morphometry features always showed excellent performance in predicting AD progression. However, the major limitation of these works was that they failed to visualize the disease-related regions on the surfaces. In the current work, our proposed FMFS model can well select the features with stronger predictive power and further visualize the ROIs on the surfaces. The proposed method is general and may be applied to analyze any general brain imaging feature data. Moreover, our experimental results show that morphometric markers from the hippocampal subiculum and CA1 subfield are apparently associated with Aβ/tau markers in all the clinically defined stages of AD and, as AD pathology progress, the ROIs showing associations are more focal. With two prediction experiments, we further demonstrate that the morphometric features on our identified ROIs show a stronger predictive power in predicting MMSE scores and future clinical decline in MCI patients. All the results indicate that FMFS is a useful screening tool to reveal associations between Aβ/tau status and hippocampal morphology across the clinically normal to dementia spectrum. Aβ/tau-associated features on ROIs could be used as potential biomarkers for the Aβ/tau pathology, perhaps as a screening tool prior to using more expensive and invasive PET techniques.

### 4.1 Aβ/Tau Associated Hippocampal Morphometry

Aβ and tau proteinopathies accelerate hippocampal atrophy leading to AD on MRI scans (Hanko et al., 2019; Maass et al., 2017; Wang et al., 2021). However, the influence of Aβ/tau deposition on hippocampal morphology in pathophysiological progression of AD is still not well understood. Some prior works (Adler et al., 2018; Shi et al., 2013a; Tsao et al., 2017) demonstrated that CA1 and the subiculum are the ROIs with the greatest abnormalities in the early stages of the AD pathophysiological process. Besides, the study of (Hanko et al., 2019) reported a significant association between tau burden and atrophy in specific hippocampal ROIs (CA1 and the subiculum), but detected no Aβ-associated hippocampal ROIs in the 42 subjects they studied.

Our work applies two kinds of morphometry measures (RD and TBM) and the novel FMFS framework to two datasets to study fine-scale morphometric correlates of Aβ and tau deposition. Both results are consistent with the prior studies noted above. Besides, we also studied the influence of Aβ/tau burden on hippocampal morphometry at different stages of AD. As the results show in **Figure 3-5**, Aβ/tau associated hippocampal ROIs are more focal as AD pathology progresses, especially at the final stage of AD itself.

### 4.2 Predictive Power of the Features on ROIs

To verify the clinical value of these identified ROIs, we compared their prediction performances to global hippocampal morphometry and Aβ/tau measures using three different machine learning models. As shown in **Table 2**, the features on our identified ROIs have superior performance for predicting clinical scores, which followed our initial hypothesis. Compared to randomly selected features, the features on ROIs show stronger predictive power, which illustrates the promise of our Federated Morphometry Feature Selection model. Additionally, these Aβ/tau-associated features always performed better than the measurements of Aβ/tau, and could be used as a potential biomarker for Aβ/tau pathology, especially as a screening indicator.

In addition, the results in **Section 3.4** further proved the stronger predictive power of the ROIs in survival analysis (of conversion from MCI to AD). Here, the univariate biomarker computed from our ROIs had better performance than the traditional hippocampal volume, which suggested the potential ability of our ROIs to study Alzheimer’s disease as a univariate biomarker. Consequently, both experiments demonstrated the effectiveness of the FMFS model.

### 4.3 Stability Analysis

To test whether the performance of our FMFS model could be affected by kinds of data distribution across institutions, we performed 5-fold cross-validation on the dataset for the study of Aβ under three conditions, including a data-centralized condition and data distributed across three institutions and five institutions. For each training data set, we randomly assigned the subjects to three institutions, five institutions, or a single institution. Besides, we also compared the performance of FMFS with DADMM and FBCD under the five-institution conditions. We perform cross-validation a total of 10 times with a sequence of regularization parameters, 1, 0.5, and 0.1, and with all the other experimental set-ups being the same as in the previous experiment. The average RMSE for the prediction of MMSE was employed to evaluate the prediction accuracy during training and testing, as shown in **Table 5**. The results indicated that different kinds of institutional distributions did not strongly influence our FMFS model.

**Table 5.** Stability analysis of FMFS across different institutional settings.

| | $\lambda$ | FMFS (1) | FMFS (3) | FMFS (5) | FBCD (5) | DADMM (5) |
| --- | --- | --- | --- | --- | --- | --- |
| Train | 1.0 | 2.80 | 2.80 | 2.80 | 2.80 | 2.80 |
|  | 0.5 | 2.70 | 2.70 | 2.70 | 2.70 | 2.70 |
|  | 0.1 | 2.43 | 2.43 | 2.43 | 2.43 | 2.44 |
| Test | 1.0 | 2.79 | 2.79 | 2.79 | 2.79 | 2.79 |
|  | 0.5 | 2.71 | 2.71 | 2.71 | 2.71 | 2.71 |
|  | 0.1 | 2.60 | 2.60 | 2.60 | 2.60 | 2.61 |

### 4.3 Limitations and Future Work

Despite the promising results are obtained by applying FMFS, there are two important caveats. First, this work is based on cross-sectional data. It would also be valuable to track the longitudinal hippocampal ROIs deformity as Aβ/tau change over time. In the future, we plan to conduct longitudinal association analyses of hippocampal features and their relation to Aβ/tau burden. Second, this work only studied the hippocampal structures, but other structures, such as the ventricles, and cortical surface metrics such as gray matter thickness or volume (Chou et al., 2009; Doherty et al., 2015) are also affected by proteinopathies. In the future, we will explore more Aβ/tau-associated brain regional abnormalities. This future work will help shed new light on the relationship of component biological processes in AD.

## 5. CONCLUSION

This work proposes a novel high-dimensional federated feature selection framework, FMFS, to study the Aβ/tau burden associated with abnormalities in hippocampal subregions on two datasets. Experimental results showed that FMFS encoded hippocampal features at different clinical stages that were associated with Aβ/tau burden. As the clinical symptoms worsen, these ROIs appear to be more focal. Our novel proposed framework achieved superior performance in efficiency compared to a similar feature selection method. To the best of our knowledge, this is the first feature selection model to study hippocampal morphometric changes with Aβ/tau burden across the AD spectrum. More importantly, this model can visualize brain structural abnormalities affected by AD proteinopathies. Beyond brain MRI, our framework may also be applied to any other kinds of medical data for feature selection.

## ACKNOWLEDGMENTS

Algorithm development and image analysis for this study were partially supported by the National Institute on Aging (RF1AG051710, R21AG065942, U01AG068057, R01AG031581, and P30AG19610), the National Institute of Biomedical Imaging and Bioengineering (R01EB025032), and the Arizona Alzheimer Consortium.

Data collection and sharing for this project was funded by the Alzheimer’s Disease Neuroimaging Initiative (ADNI) (National Institutes of Health Grant U01 AG024904) and DoD ADNI (Department of Defense award number W81XWH-12-2-0012). ADNI is funded by the National Institute on Aging, the National Institute of Biomedical Imaging and Bioengineering, and through generous contributions from the following: Alzheimer’s Association; Alzheimer’s Drug Discovery Foundation; BioClinica, Inc.; Biogen Idec Inc.; Bristol-Myers Squibb Company; Eisai Inc.; Elan Pharmaceuticals, Inc.; Eli Lilly and Company; F. Hoffmann-La Roche Ltd and its affiliated company Genentech, Inc.; GE Healthcare; Innogenetics, N.V.; IXICO Ltd.; Janssen Alzheimer Immunotherapy Research & Development, LLC.; Johnson & Johnson Pharmaceutical Research & Development LLC.; Medpace, Inc.; Merck & Co., Inc.; Meso Scale Diagnostics, LLC.; NeuroRx Research; Novartis Pharmaceuticals Corporation; Pfizer Inc.; Piramal Imaging; Servier; Synarc Inc.; and Takeda Pharmaceutical Company. The Canadian Institutes of Health Research is providing funds to support ADNI clinical sites in Canada. Private sector contributions are facilitated by the Foundation for the National Institutes of Health (www.fnih.org). The grantee organization is the Northern California Institute for Research and Education, and the study is coordinated by the Alzheimer’s Disease Cooperative Study at the University of California, San Diego. ADNI data are disseminated by the Laboratory for Neuro Imaging at the University of Southern California.

Data used in the preparation of this article were obtained from the Alzheimer’s Disease Neuroimaging Initiative (ADNI) database (adni.loni.usc.edu). As such, the investigators within the ADNI contributed to the design and implementation of ADNI and/or provided data but did not participate in the analysis or writing of this report. A complete listing of ADNI investigators can be found at: http://adni.loni.usc.edu/wp-content/uploads/how_to_apply/ADNI_Acknowledgement_List.pdf.

